# Reactivation of developmentally silenced globin genes through genomic deletions reveals that enhancer distance matters

**DOI:** 10.1101/2025.01.13.632719

**Authors:** Anna-Karina Felder, Sjoerd J.D. Tjalsma, Han J.M.P. Verhagen, Rezin Majied, Marjon J.A.M. Verstegen, Thijs C.J. Verheul, Rebecca Mohnani, Richard Gremmen, Peter H.L. Krijger, Sjaak Philipsen, Emile van den Akker, Wouter de Laat

## Abstract

The human genome contains regulatory DNA elements, enhancers, that can activate gene transcription over long chromosomal distances. Here, we show that enhancer distance can be critical for gene silencing. We demonstrate that linear recruitment of the normally distal *HBB* super-enhancer to developmentally silenced *HBG* promoters, through deletion or inversion of intervening DNA sequences, results in strongly reactivated *HBG* expression in adult erythroid cells and *ex vivo* differentiated hematopoietic stem and progenitor cells. A similar observation is made in the *HBA* locus, where deletion-to-recruit of the distal enhancer strongly reactivates embryonic *HBZ* expression. Overall, our work assigns function to seemingly non-regulatory genomic segments: by providing linear separation they may support genes to autonomously control their transcriptional response to distal enhancers.

## Introduction

The vast majority (98%) of the human genome consists of non-coding DNA sequences. They provide a home for over a million of transcriptional regulatory elements that collectively coordinate the cell-specific gene expression programs driving cell differentiation, embryonic development and organogenesis (1, 2). Developmental gene activation depends on *cis*-acting DNA elements called enhancers. Enhancer action requires functional communication between transcription (co)factors recruited to the enhancer and the gene promoter. This functional communication is thought to occur through chromatin looping, which brings distant regulatory elements into close physical proximity to their target promoters (3, 4). Endogenous enhancers can locate anywhere between 0kb to 1500kb away from their designated target gene (5), raising the question whether linear distance has any functional implication. Random integration of a reporter gene in mice indicated that the likeliness of being expressed in a given tissue correlated with linear proximity to an enhancer active in that tissue (6). Transcript levels generally decreased with increasing enhancer-promoter distances (7–10). Artificial regulatory landscapes, in which a reporter gene was activated by an enhancer at varying distances (0-400kb), corroborated that enhancer distance anti-correlated with transcriptional activity and stability: the more distal the enhancer was positioned, the higher the proportion of cells silencing the reporter. Deletion of enhancer-promoter intervening sequences in the silenced cells resulted in strong transcriptional reactivation of the reporter (8). This suggested that by keeping enhancers at a distance, genes may gain control over their response to enhancers (8). Here, we aimed to study the role of genomic enhancer-promoter distance at the endogenous human α-globin (*HBA*) and β-globin (*HBB*) locus.

The *HBA* locus contains an embryonic gene, *HBZ*, and two adult genes, *HBA1* and *HBA2*. Their expression is regulated by distal enhancers that are active in erythroid cells throughout development (11, 12). The *HBB* locus consists of five β -like globin genes: the embryonic *HBE* gene precedes two fetal *HBG* genes, which are followed downstream by the adult *HBD* and *HBB* genes. Their activation and expression levels are controlled by the upstream Locus Control Region (LCR) (13), a super enhancer that is active at all stages of erythroid development.

Silencing is intrinsically controlled by the promoter regions of the developmentally expressed globin genes (14). For instance, *HBG* inactivation is the consequence of repressor proteins recruited to the *HBG* gene promoters, with BCL11A binding 115bp upstream the TSS being the best characterized (15, 16). Disruption of this binding site in the *HBG* gene promoter leads to *HBG* reactivation (17, 18). Upstream of the *HBG* gene promoters, in the 25 kb interval that separates *HBG* from the LCR, there are no indications for regulatory sequences preventing LCR-mediated activation of the *HBG* promoters in adult erythroid cells. Interestingly, HBG repression in adult red blood cell precursors can also be reversed by forced looping of the LCR to the HBG promoters (19). The current idea therefore is that stage-specific recruitment of repressor proteins to the *HBG* promoters prevents LCR engagement with the *HBG* genes, resulting in developmental silencing.

We hypothesized that developmentally silenced genes may be reactivated upon forced linear recruitment of enhancers through CRISPR-Cas deletion of intervening sequences: delete-to-recruit (Del2Rec). We tested this by applying Del2Rec to the globin genes in human cells.

## Results

### Forced linear proximity of the *HBB* super-enhancer reactivates *HBG* expression

The *HBB* LCR spans nearly 20 kb and has five hypersensitive sites (HS1-5) (Figure 1A), with different intrinsic enhancer capacities(27, 28). HS2, -3 and -4 are all potent individual enhancers, while HS1 and -5 lack strong intrinsic enhancer activity. To develop Del2Rec, we therefore designed guide RNAs (gRNAs) close to HS2. The accompanying gRNA was placed upstream of the *HBG2* promoter. Combined, they generate a 25kb deletion placing HS5-4-3-2 immediately upstream of the *HBG2* promoter, leaving all reported repressor protein binding sites intact (Figure 1A). Several gRNAs near HS2 and *HBG2* were tested for efficiency. Based on TIDE analysis (29), we selected gRNA-HS2 (located inside HS2) and gRNA-500 (located 500 bp upstream of the *HBG2* TSS); gRNA-500 is unique for the *HBG2* promoter and leaves the *HBG1* promoter intact.

**Figure 1.**
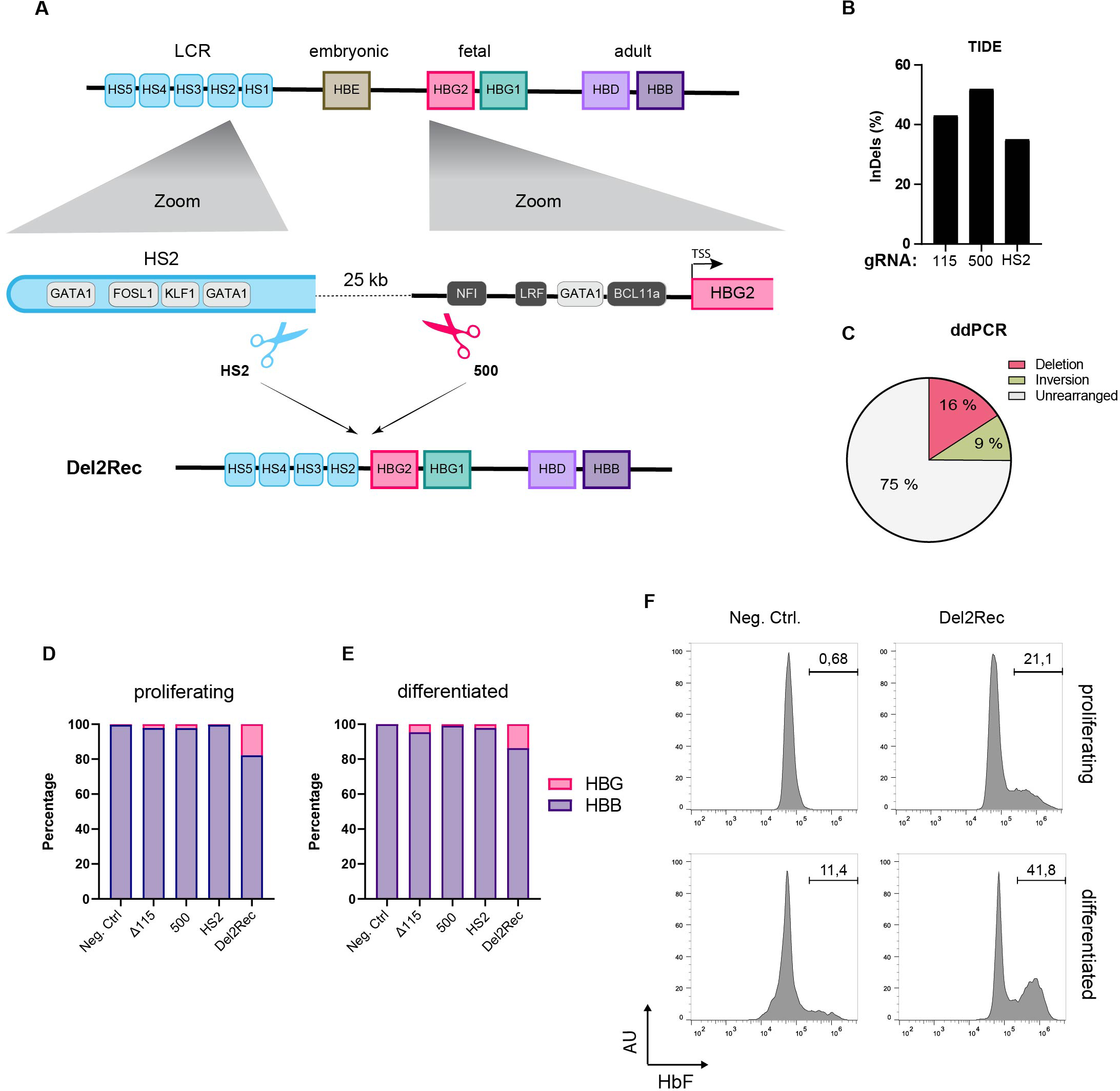
Linear recruitment of the LCR reactivates fetal globin in HUDEP-2 cells. (A) Schematic representation of the *HBB* locus on chromosome 11, depicting the hypersensitive sites of the locus control region (light blue), *HBE* (brown), *HBG2* (pink), *HBG1* (dark mint), *HBD* (lilac), and *HBB* (purple) genes. Below (left) zoom in on HS2 showing the most relevant TF-BS (grey) and binding site of gRNA-HS2. On the right side a zoom-in on the *HBG2* promoter with the most relevant TF-BS (gray) and position of gRNA_500. At the bottom the 25kb deletion (Del2Rec) generated by combination of gRNA_500 and gRNA-HS2. (B) Frequency of InDels, measured by TIDE analysis for Δ115, 500 and HS2 gRNAs in single-gRNA transfected samples. (C) Frequency of the 25-kb deletion (raspberry), inversion (olive) and unmodified (light gray), measured by ddPCR in Del2Rec condition. (D) RT-qPCR analysis of HBB and HBG mRNA levels in proliferating (day 0) cells. HBB and HBG mRNA expression was normalized to actin mRNA and is displayed as percentage of HBB plus HBG. (E) Same as in (D) but for cells differentiated for 10 days. (F) Flow cytometry plots showing the percentage of HbF-positive cells in proliferating (top row) and after 10 days of differentiation (bottom row) cells.

We first assayed the transcriptional consequences of Del2Rec in HUDEP-2 cells, a model system for human adult erythroid cells (21). TIDE analysis(29) revealed that gRNA-HS2 and gRNA-500, when individually transfected as Cas9 RNPs into HUDEP-2 cells, reached local insertion/deletion (indel) rates of respectively 35% and 52% (Figure 1B). This was similar to the 43% local indel rate measured for gRNA-115 (30) that served as a control. We applied digital droplet PCR (ddPCR) (Figure S1A) to quantify the 25 kb deletion, inversion, and unrearranged allele frequencies upon nucleofection of gRNA-HS2 + gRNA-500 RNPs. This showed that 16% of the alleles carried a deletion and 9% an inversion of the 25 kb fragment (Figure 1C).

We then tested whether Del2Rec caused *HBG* gene reactivation. We measured *HBG* and *HBB* expression levels by RT-qPCR. Despite the fact that only 16% of the alleles carried the 25 kb deletion, we observed considerable *HBG* gene reactivation before and after differentiation (Figure 1D-E). Consistent with these data, flow cytometry revealed that Del2Rec increased the proportion of cells with high fetal hemoglobin levels (F-cells; Figure 1F). Replicate experiments confirmed that Del2Rec induced *HBG* gene reactivation (Figure S2). Together, these results indicate that forced linear proximity of the LCR can overwrite the promoter-encoded developmental silencing of the *HBG* genes in adult red blood cell precursors.

### Del2Rec drives activation of the proximal *HBG2* gene and suppression of the distal *HBB* gene

To obtain direct evidence that Del2Rec leads to *HBG* gene reactivation we generated clonal HUDEP-2 cell lines carrying the intended 25 kb deletion on both alleles. These clonal cell lines, Del2Rec-1 to -3, carried alleles with slightly different deletion junctions. All alleles retained at least 493 bp of the *HBG2* promoter and the complete *HBG1* gene. For comparison, we applied gRNA1617 (24) which targets the erythroid-specific enhancer of *BCL11A* (hereafter referred to as ΔBCL11A). In two clonal HUDEP-2 lines (ΔBCL11A-1 and ΔBCL11A-2), with indels on both alleles of the *BCL11A* enhancer, ATAC-seq showed that the *BCL11A* enhancer was no longer accessible, and BCL11A protein was no longer detectable by western blot (Figure S3A-B). In addition, we used gRNA-115 (30) to generate two HUDEP-2 clonal lines (Δ115-1 and Δ115-2) with homozygous disruption of the BCL11A binding sites in the *HBG* promoters (Figure S3C).

The Del2Rec lines showed strong *HBG* upregulation in proliferation conditions (Figure 2A). Upon differentiation, we found further increased *HBG* transcript levels (Figure 2A). Concomitantly *HBB* transcript levels increased as well, but not to control HUDEP-2 cells levels (Figure 2B), suggesting that the activated *HBG* genes compete with the more distal *HBB* genes for activation by the LCR (31–33). Next, we determined whether the proximal or distal *HBG* gene was activated by the LCR. Using a *HBG* gene-specific RT-qPCR strategy, we observed that the Del2Rec clones nearly exclusively reactivated the proximal *HBG2* gene (Figure 2C-D). In contrast, the Δ115 and ΔBCL11A lines expressed *HBG2* and *HBG1* at similar levels (Figure 2C-D). Overall *HBG* expression levels in the Del2Rec lines were similar to those measured in the Δ115 lines and higher than found in the ΔBCL11A lines. To exclude that elevated *HBG2* transcript levels were the consequence of LCR-initiated enhancer transcripts reading into *HBG2*, we analyzed the transcripts with different PCR amplicons covering the promoter area and the *HBG2* TSS. This revealed that the elevated transcript levels were not LCR-initiated read-through events, but bona fide gene transcripts initiated from the *HBG2* TSS (Figure 2E).

**Figure 2.**
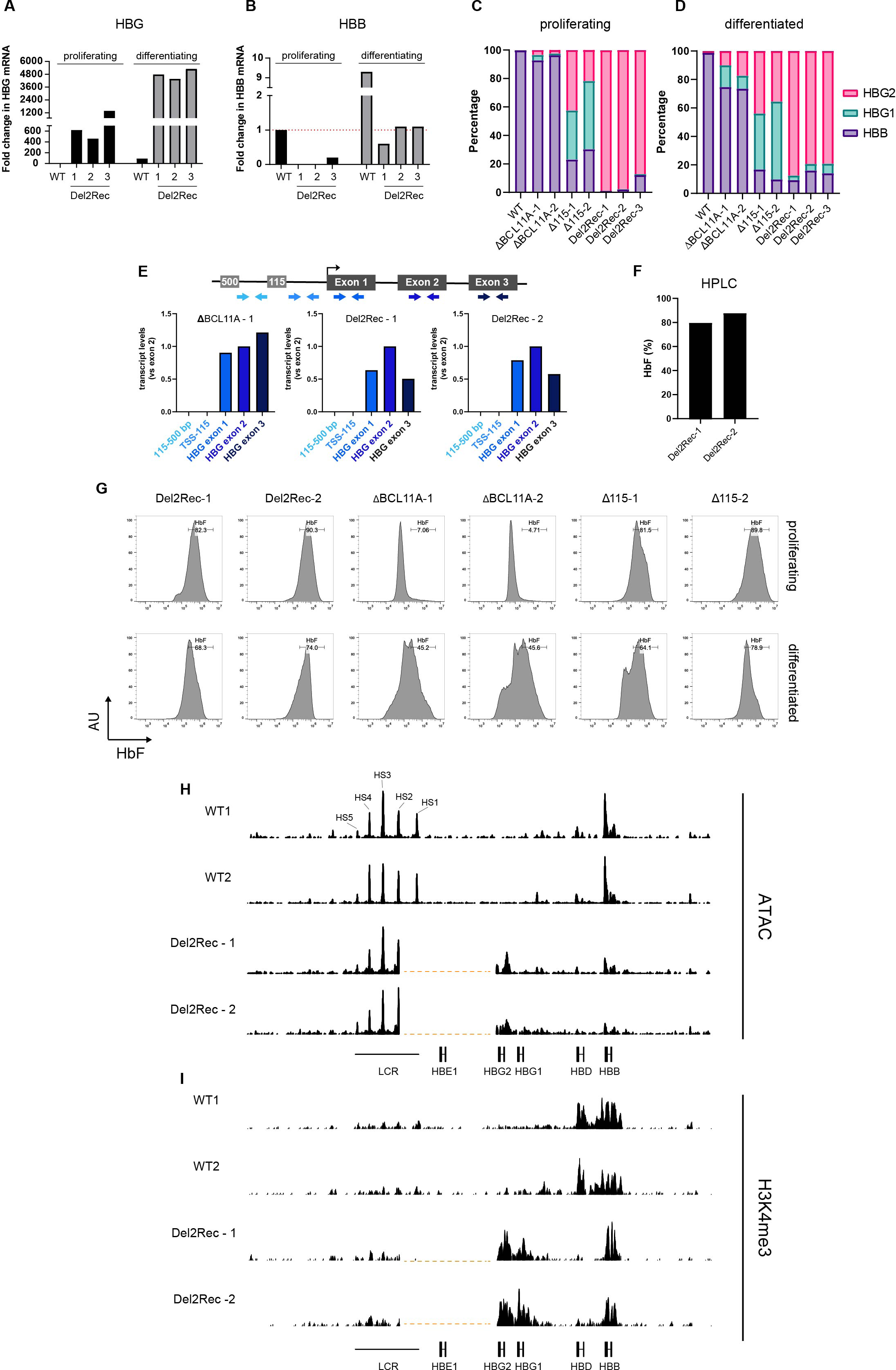
Del2Rec clonal HUDEP-2 cell lines give insight into *HBG* reactivation mechanism. (A) RT-qPCR analysis of HBG mRNA levels in proliferating (left, black bars) and 10 days differentiated (right, black bars) clonal cell lines. Expression was normalized to actin mRNA and fold-change was calculated over proliferating control cells. (B) Same as in (A) but fold-change of HBB mRNA. (C) RT-qPCR analysis of HBB, HBG1 and HBG2 mRNA levels in proliferating clonal cell lines. HBB, HBG1 and HBG2 mRNA expression was normalized to actin and displayed as percentage of HBB + HBG1 + HBG2. (D) Same as (C) but cells differentiated for 10 days. (E) Schematic representation of primers used in RT-qPCR analysis of HBG transcripts (top). Fold-change normalized for primer efficiency based on DNA amplification with HBG2 exon 2 set to 1, shown in arbitrary units of a ΔBCL11A clone and two Del2Rec deletion clones (bottom). (F) HbF of two clonal Del2Rec deletion cell lines measured by HPLC. Percentage of HbF calculated over the total Hb tetramers. (G) Representative flow cytometry plots showing the percentage of HbF-positive cells in proliferating clonal cell lines (top row) and after 10 days of differentiation (bottom row). (H) ATAC–seq tracks of the *HBB* gene cluster in differentiated (10 days) WT HUDEP-2 cells and two Del2Rec clones. Vertical dashed lines indicate deletion generated in Del2Rec clones. Signal is scaled to genome wide TSS signal. Max scales were set to highest scaled signal in *HBB* locus. (I) H3K4me3 CUT&RUN tracks of the *HBB* gene cluster in two differentiated WT HUDEP-2 samples and two Del2Rec clones. Vertical dashed lines indicate deletion generated in Del2Rec clones. Scaling is based on average coverage at *HBA* locus.

For the Del2Rec lines, HPLC showed that 80-90% of hemoglobin consisted of fetal hemoglobin (HbF; Figure 2F), and by flow cytometry an almost pure population of F cells was observed (Figure 2G). This was comparable to HbF and F cell levels observed in Δ115 cells. In the ΔBCL11A lines, high HbF and F cell levels were only seen after differentiation. In the Del2Rec lines, ATAC-seq (Figure 2H) and CUT&RUN for H3K4me3 (Figure 2I) demonstrated that linear recruitment of the LCR led to chromatin opening and H3K4 trimethylation of the *HBG2* promoter. Interestingly, ATAC-seq indicated partial reduction in accessibility of the more distal adult *HBB* gene (Figure 2H), likely as a consequence of Del2Rec-induced gene competition for activation by the LCR.

Finally, we investigated *HBG* transcript and protein levels in two clonal cell lines with monoallelic 25 kb deletion and an intact wildtype allele. While *HBG* transcript levels were reduced compared to those in the homozygous Del2Rec cell lines, still 60-70% of the β-globin-like transcripts were of *HBG* origin (Figure S4A-B) and by flow cytometry a high percentage of F cells was observed (Figure S4C). Thus, Del2Rec can support effective *HBG2* reactivation upon monoallelic *HBB* locus rearrangement.

We conclude that Del2Rec causes strong reactivation of *HBG2* expression in adult erythroid cells. We therefore propose that linear distance between the LCR and the genes is essential for the *HBG* promoters to autonomously control their developmental silencing in adult erythroid cells.

### Forced linear proximity of the enhancer can overwrite *HBG* silencing in an SCD model and CD34+ HSPCs differentiated to the erythroid lineage

Partially restored *HBG* expression can ameliorate the symptoms of sickle cell disease (SCD) and β-thalassemia (β-thal)(34–36), two common severe monogenic diseases caused by *HBB* variants. We therefore tested whether Del2Rec would also be effective in two erythroid progenitor cell lines (SCD1 and SCD2) derived from SCD patients. TIDE analysis showed that Del2Rec nucleofection was relatively efficient (60-80%), compared to control nucleofections with gRNA-115 (15-55%) and the BCL11A targeting gRNA1617 (55-60%) (Figure 3A). ddPCR showed that Del2Rec treatment resulted in 24%/22% (SCD1) and 35%/12% (SCD2) alleles with the 25 kb deletion/inversion, respectively (Figure 3B).

**Figure 3.**
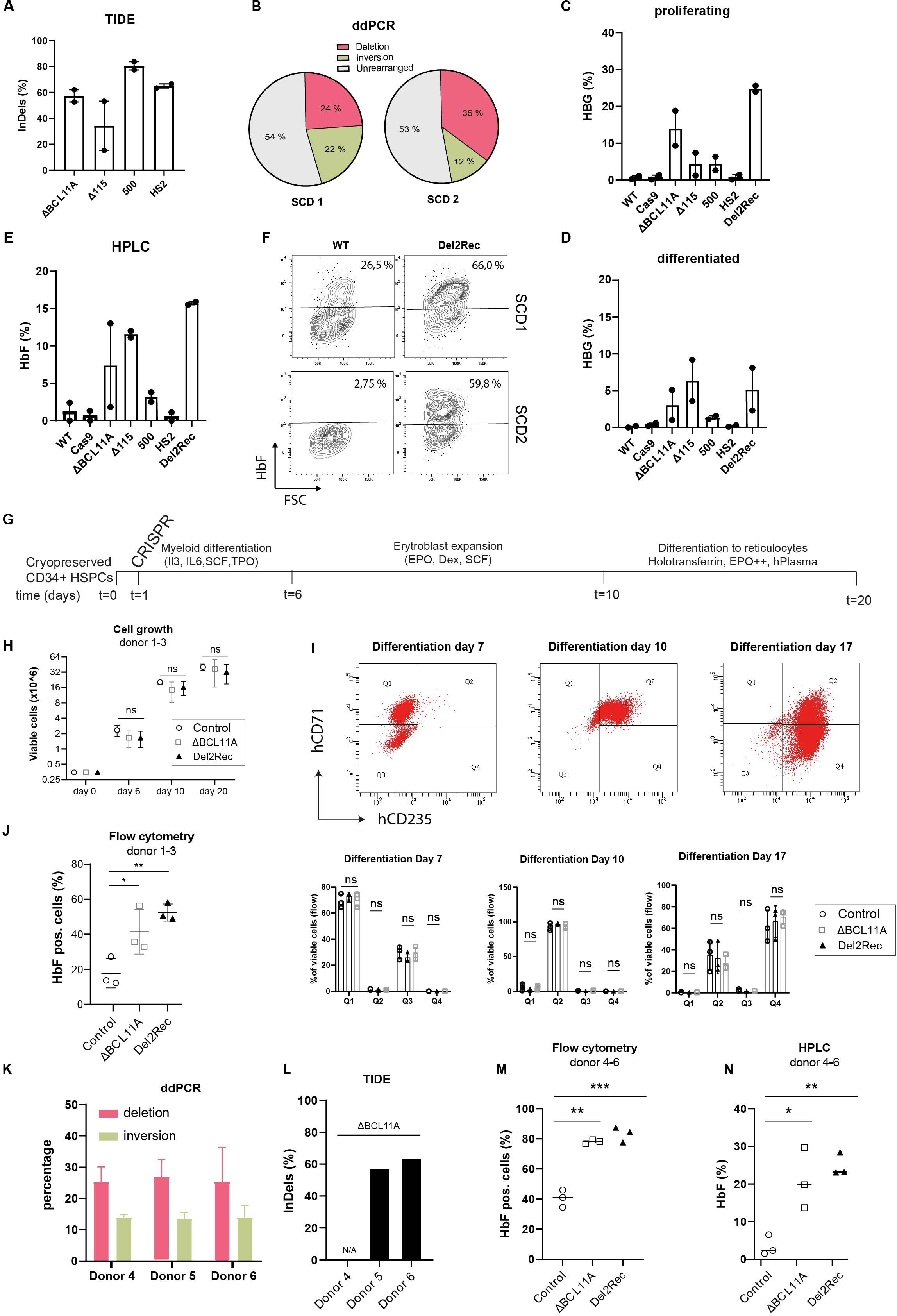
Linear recruitment of the LCR reactivates fetal globin in immortalized, patient-SCDs cell lines and HSPCs. (A) Frequency of InDels, measured by TIDE analysis for ΔBCL11A (gRNA1617), Δ115, 500 and HS2 gRNAs in single-gRNA transfected samples in two SCD cell lines shown as average ± SEM. (B) Frequency of the 25-kb deletion (raspberry), inversion (olive) and unmodified (light gray), measured by ddPCR in Del2Rec cells. (C) RT-qPCR analysis HBG mRNA levels in proliferating cells, displayed as the average ± SEM in percentage of total HBB+HBG mRNA. (D) Same as in c) but for cells differentiated for 4 days. (E) HbF of two REB-SCD cell lines measured by HPLC shown as average ± SEM. Calculated as percentage of HbF over the total Hb tetramers. (F) Flow cytometry plots showing the percentage of HbF positive cells in two REB-SCD cell lines. Note that SCD1 shows relatively high baseline HbF levels. (G) Schematic overview of culture conditions of CD34+ HSPCs. (H) Cell counts of edited HSPCs of donor 1-3 (day 0) differentiating towards reticulocytes (day 20). Dots show the mean cell counts and error bars show the SD. Control is untreated donor HSPCs. (I) Flow cytometry plots show representative differentiation patterns of HSPCs of donors 1-3 to reticulocytes measured on indicated time points (top). Graphs show the percentage of indicated cell populations for each individual donor. Error bars show the SD of a triplicate (bottom). (J) Graph showing percentage of HbF positive cells of donors 1-3 as measured by flow cytometry. Error bars show the SD of a triplicate. (K) Frequency of the 25-kb deletion (raspberry) and inversion (olive) measured by ddPCR on Del2Rec transfected cells. Values were calculated as an average of two measurements from gDNA isolated from donors 4-6. Error bars represent SD. (L) Frequency of InDels, measured by TIDE analysis for ΔBCL11A (gRNA1617) of donors 4-6. (M) Percentage of F-cells plotted as averages of 3 donors after 12 days of differentiation. An one-way ANOVA and Tukey’s post-hoc test was used to calculated statistics. ns = not significant, * = p< 0.05 , ** p<0.01 , ***p<0.001. (N) HbF of donors 3-6 measured by HPLC 3 days after differentiation in control, ΔBCL11A and Del2Rec conditions. We calculated the percentage of HbF over the total Hb tetramers.

We examined *HBG* versus *HBB* transcript ratios in the edited cells by RT-qPCR. While the individual gRNAs HS2 and -500 gave little *HBG* reactivation, both cell lines strongly responded to Del2Rec (Figure 3C-D). This was confirmed at the protein level by HPLC (Figure 3E) and by flow cytometry for F cells (Fig 3F and Figure S5). We conclude that Del2Rec also efficiently induces *HBG* reactivation in erythroid progenitor cell lines derived from SCD patients.

Thus far, we induced deletions in cells already committed to the erythroid lineage, i.e. having an open *HBB* locus with an accessible LCR primed to engage with downstream accessible β-like globin genes. Therefore, we next determined the effects of Del2Rec in primary CD34+ hematopoietic stem and progenitor cells (HSPCs). We edited CD34+ HSPCs, isolated from three healthy donors (Figure 3G); gRNA1617 (24), targeting the BCL11A enhancer, was used as a positive control. The proliferation, erythroid specification and differentiation characteristics of Del2Rec-edited HSPCs were similar to those of control cells (Figure 3H-I).

Importantly, Del2Rec resulted in high *HBG* activation, at levels comparable to those obtained with the BCL11A enhancer gRNA (Figure 3J).

To further investigate this, we repeated the experiment with 3 new donors and used ddPCR to quantify Del2Rec edits (Figure 3K). This revealed that Del2Rec induced deletions/inversions in, on average, 26%/14% of the alleles respectively (Figure 3K). TIDE analysis showed that gRNA1617 modified ∼60% of the *BCL11A* alleles (Figure 3L), as reported before (24). Upon differentiation, flow cytometry showed that both approaches yielded around 80% F-cells as compared to 35-45 % in the controls (Figure 3M). HPLC analysis revealed that cells expressed on average 20-25% HbF with both approaches, but Del2Rec appeared to give less variable expression, as compared to targeting of the *BCL11A* enhancer (Figure 3N). Collectively, we conclude that Del2Rec can reactivate *HBG* expression in erythroid cells derived from CD34+ HSPCs.

### Forced linear proximity through inversions also drives *HBG* reactivation

A relatively high proportion of alleles showed inversion of the 25 kb fragment upon Del2Rec. We explored this to further test our hypothesis that *HBG* activation can be induced by forced linear proximity of enhancers. For this, we designed two alternative editing strategies. The first used gRNA-500 combined with a novel gRNA-HS2/3 targeting the region between HS2 and HS3. gRNA-HS2/3 splits the LCR between HS5-4-3 and HS2-1. By combining gRNA-500 and gRNA-HS2/3, the resulting deletion would place *HBG2* immediately downstream of HS5-4-3, while the inversion would place the gene downstream of the inverted HS2-1 (Figure 4A). Editing HUDEP-2 cells with gRNA-500 + gRNA-HS2/3 in bulk resulted in the anticipated deletions/inversions (Figure S6A) and induced high levels of *HBG* expression at the RNA (Figure 4B) and protein level (Figure 4C). Furthermore, it was predominantly *HBG2* that was reactivated, not *HBG1* (Figure 4B). To test whether inversion events contribute to reactivated *HBG2* expression, we isolated clonal cell lines carrying the 25 kb genomic interval inverted on one of the alleles. Four clonal lines were selected (Figure S6B) and they all showed strong *HBG* reactivation (Figure 4D). Thus, inversions can also cause reactivation of developmentally silenced genes, provided that they recruit a sufficiently strong enhancer in close proximity to the to-be-activated gene. Importantly, in these inversion clones strong *HBG* reactivation occurs in the presence of all sequences of the *HBB* locus. This is further evidence that developmental silencing of *HBG* expression is primarily controlled by the *HBG* promoters.

**Figure 4.**
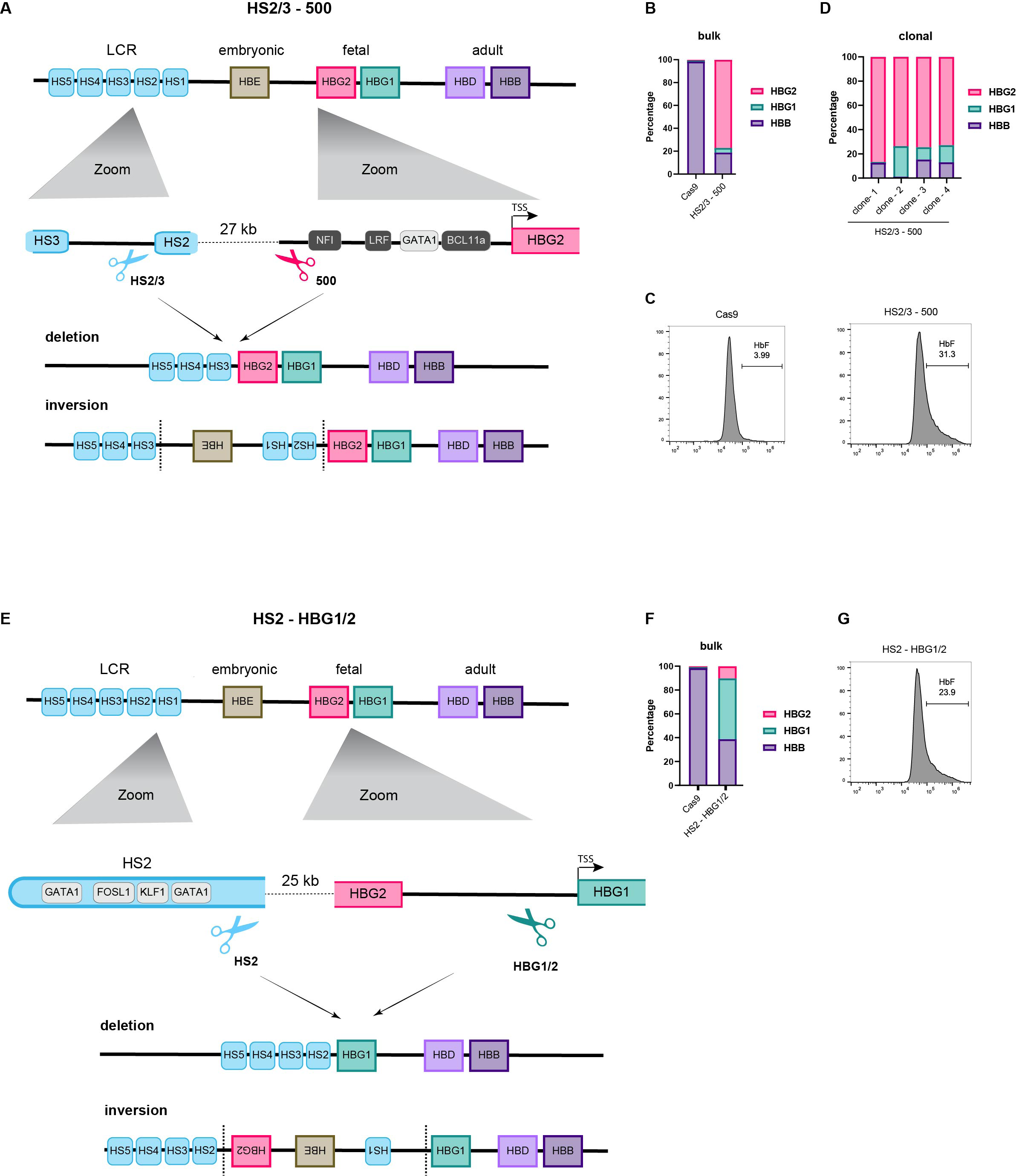
Inversion of intervening sequence can also reactivate silenced genes. (A) Schematic representation of the *HBB* locus on chromosome 11, depicting the hypersensitive sites of the locus control region (light blue), *HBE* (brown), *HBG2* (pink), *HBG1* (dark mint), *HBD* (lilac), and *HBB* (purple) genes. Below (left) zoom in on HS2 and HS3, showing the position of gRNA-HS2/3. On the right side a zoom in on the *HBG2* promoter with the most relevant TF-BS (gray) and position of gRNA_500. At the bottom the 27kb deletion generated by combination of gRNA_500 and gRNA-HS2/3. Below a representation of the inversion generated by the same gRNAs. (B) RT-qPCR analysis of HBB, HBG1 and HBG2 mRNA levels in proliferating bulk cell populations. HBB, HBG1 and HBG2 mRNA expression was normalized to actin and displayed as percentage of HBB + HBG1 + HBG2. WT is the parental lentiviral Cas9-expressing cell line without gRNAs. (C) Flow cytometry plots showing the percentage of HbF-positive cells in proliferating bulk cell populations. (D) RT-qPCR analysis of HBB, HBG1 and HBG2 mRNA levels in four proliferating heterozygous inversion clones. HBB, HBG1 and HBG2 mRNA expression was normalized to actin and displayed as percentage of HBB + HBG1 + HBG2. (E) Same as (A) but a schematic representation of gRNAs HS2 and HBG1/2, cutting between HBG1 and HBG2. (F) RT-qPCR analysis of HBB, HBG1 and HBG2 mRNA levels in proliferating bulk cell populations. HBB, HBG1 and HBG2 mRNA expression was normalized to actin and displayed as percentage of HBB + HBG1 + HBG2. WT as in (B). (G) Flow cytometry plot showing the percentage of HbF-positive cells in proliferating bulk cell population.

To further substantiate this notion, we combined gRNA-HS2 with new gRNA-HBG1/2 that targets the region between *HBG2* and *HBG1* (Figure 4E). HUDEP-2 cells edited by gRNA-HS2 + gRNA-HBG1/2 generated the expected deletions/inversions (Figure S6A,B), and again strongly reactivated *HBG* expression (Figure 4F-G). However, as predicted from the created deletion, now expression of both *HBG1* and *HBG2* was observed. *HBG2* can only be reactivated on inverted alleles, since it is removed from deleted alleles. In such inversions, *HBG2* is reactivated despite retaining all its natural 25 kb upstream sequences, further ruling out that this genomic interval contains regulatory sequences important for developmental *HBG* silencing. The 25 kb interval performs its function by creating genomic distance between the *HBG* genes and the LCR enhancers. This linear separation enables the fetal *HBG* genes to negate the enhancers and execute their promoter-encoded developmental silencing program in adult erythroid cells.

### Del2Rec reactivates the embryonic *HBZ* gene in the *HBA* locus

Next, we investigated if the principle of reactivating developmentally silenced genes by Del2Rec could be extended to other loci. The *HBA* locus (Figure 5A) contains an embryonic gene, *HBZ*, that is completely silenced (Figure 5B), and two active adult globin genes (*HBA1* and *HBA2*) in HUDEP-2 cells. Their main enhancer, R2, is located ∼40 kb upstream of *HBZ*(*11*). We selected four gRNAs targeting sites between 245 and 622 base pairs upstream of the *HBZ* TSS(*37*), and combined each with a gRNA directly downstream of enhancer R2(38). All four gRNA combinations resulted in the expected deletions (Figure S6A) that linearly recruited R2 in close proximity to the *HBZ* promoter, and in all cases this was accompanied by an increase in *HBZ* expression (Figure 5B).

**Figure 5.**
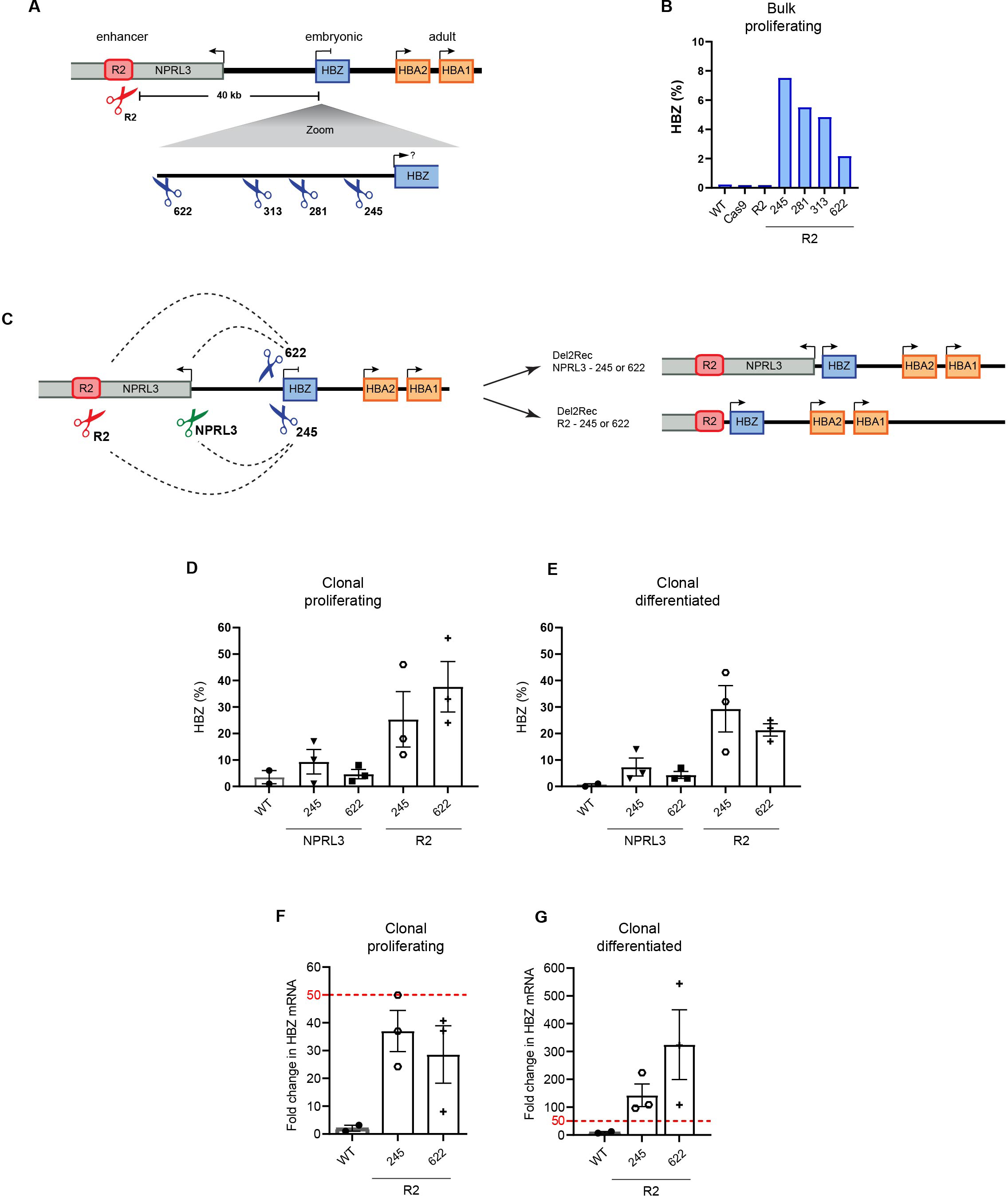
Linear recruitment of the R2 enhancer reactivates *HBZ* in HUDEP-2 cells. (A) Schematic overview of the *HBA* locus on chromosome 16, depicting R2 enhancer (red), *HBZ* (blue), *HBA1/2* (orange), and *NPRL3* (grey) genes. The red scissor displays the gRNA at the R2 enhancer. Below a zoom in on *HBZ* showing the binding sites of the four gRNAs (blue scissors) with the distance to the *HBZ* TSS. (B) RT-qPCR analysis of HBA and HBZ mRNA levels in proliferating (day 0) bulk nucleofected HUDEP-2 cells recruiting R2 to different distances from the *HBZ* TSS. HBA and HBZ mRNA expression was normalized to actin mRNA and is displayed as percentage HBZ of HBA plus HBZ. (C) Schematic representation of deletions performed in the *HBA* locus. Two gRNAs at the *HBZ* promoter were either combined with a gRNA at the R2 enhancer, or 490 bp upstream of the *NPRL3* TSS. Dotted lines represent deletions obtained upon treatment with two indicated gRNAs. (D) Same as (B), but RT-qPCR performed in heterozygous-deleted HUDEP-2 lentivirus-transduced clonal cell lines. WT describes cell lines derived in the same experiment and clonal outgrowth but genetically un-rearranged. Percentages displayed as averages ± SEM of 2-3 clones. (E) Same as (B), but for clones differentiated for three days. (F) and (G) RT-qPCR results of R2-*HBZ* deletion clones, normalized to HBZ levels in proliferating WT clones.

We isolated clonal lines with heterozygous deletions bringing the R2 enhancer either 622 bp or 245 bp upstream of the *HBZ* TSS. We also generated heterozygous clonal lines with smaller deletions of approximately 15 kb, that linked the 622 bp or 245 bp *HBZ* promoters to 490 bp in front of the *NPRL3* promoter; *NPRL3* is located in between the *HBA* genes and R2 and transcribes away from the *HBA* genes (Figure 5C, Figure S7B-C). The recruitment of the *NPRL3* promoter to either version of the *HBZ* promoter did not result in reactivation of *HBZ* expression (Figure 5D-E). In contrast, the recruitment of enhancer R2 to the two differently sized *HBZ* promoters resulted in strong reactivation of *HBZ* expression (Figure 5D-G). Similar to Del2Rec in the *HBB* locus, increased *HBZ* transcript levels were the consequence of regained promoter activity, not of enhancer transcripts extending into the *HBZ* gene (Figure S5D). Collectively, we conclude that Del2Rec can be applied to (re)activate gene expression at other loci than *HBB*.

## Discussion

Our demonstration that Del2Rec can reactivate the *HBG* genes aligns well with the previous remarkable observation that forced looping of the LCR to the *HBG* genes reactivates their expression in adult red blood cells(19). In these experiments, the complete locus, including the 25 kb stretch of intervening DNA sequences as well as the *HBG* promoters, were kept intact. *HBG* reactivation was accomplished by forcing a chromatin loop that spatially recruited the LCR to the *HBG* promoters(19). Together with our work this shows that to establish and maintain the *HBG* promoters in their silenced state in adult erythroid cells, the LCR needs to be kept away from these promoters. When brought in proximity through forced chromatin looping or by deletions/inversions, the LCR overrides the developmental silencing program of the *HBG* promoter and activates it. We show that the same principle is also operational at the *HBA* locus.

Chromosomal rearrangements resulting in enhancer hijacking and ectopic gene activation have been extensively documented, as they can drive cancer and cause developmental diseases. These hijacking events either involve *trans*-rearrangements that fuse different chromosomes to recruit enhancers to genes they would normally never engage with, or *cis*-rearrangements that eliminate the insulating boundaries of topologically associating domains that serve to prevent undesired interactions between enhancers and non-target genes (39, 40). Here, we show that forced recruitment or ‘enhancer hijacking’ can also be accomplished by the deletion or inversion of a genomic stretch of DNA without intrinsic insulating capacity, that lies in between an enhancer and gene and thus creates linear distance between these elements. Genes require this chromosomal distance to negate activation by the enhancer, thus enabling execution of the promoter-encoded developmental silencing program. This work therefore assigns a regulatory function to some of the intrinsically non-functional stretches of genomic DNA.

Understanding the contribution of enhancer distance to gene regulation provides valuable insights into therapeutic opportunities for globin gene reactivation. For α-thalassemia caused by HBA deletions, restoring HBZ levels could be a promising approach, though bringing R2 near the HBZ promoter disrupts the essential NPRL3 gene (12). In the HBB locus, reactivation of HBG, for example by targeting BCL11A, has already shown therapeutic success in SCD and β-thalassemia (34). Whether recruitment of the LCR to *HBG* may offer an orthogonal therapeutic approach requires further investigation.

## Supplementary Materials

**Fig. S1.**
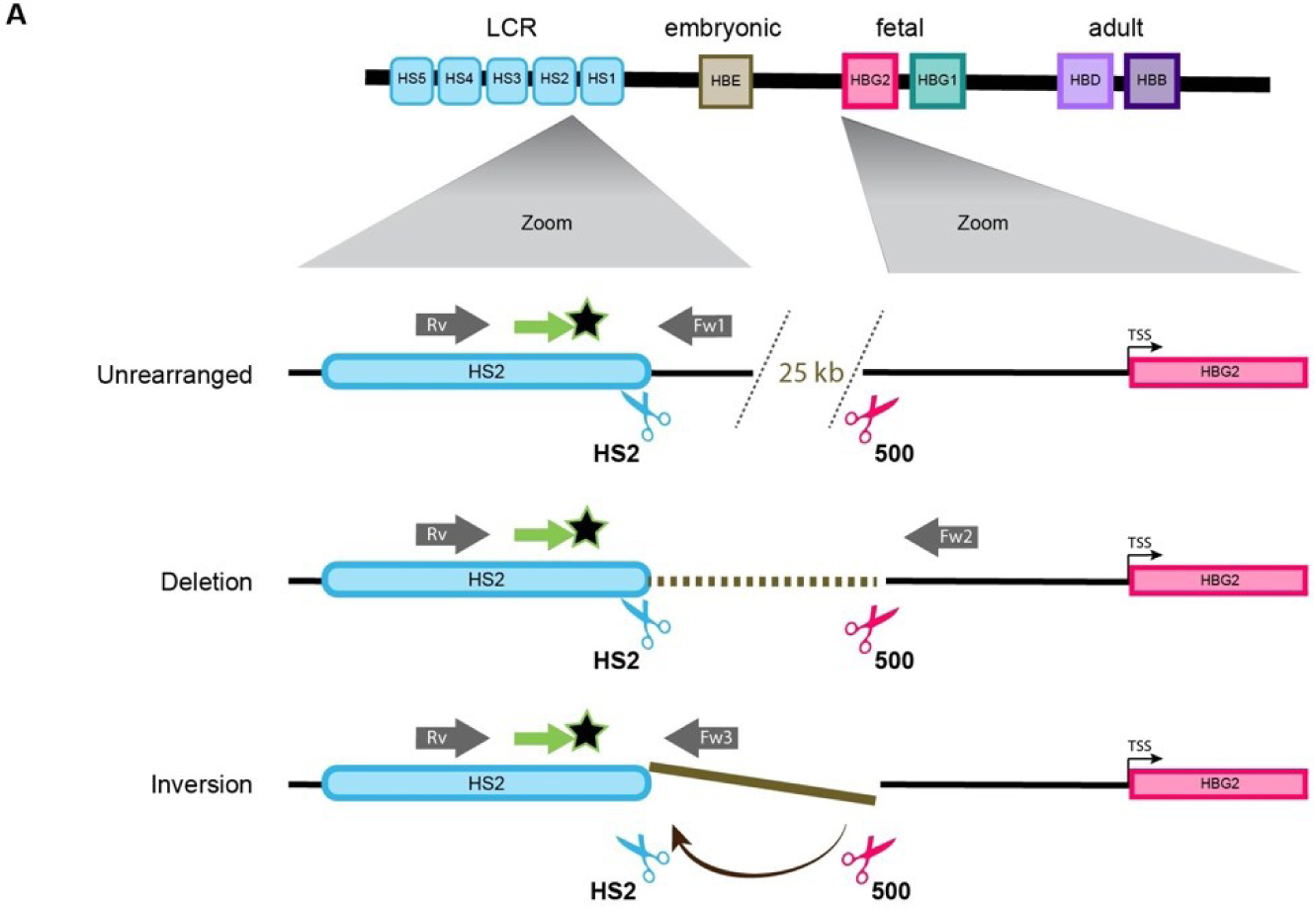
Schematic overview of ddPCR assay. a) Schematic overview of the ddPCR assay measuring the 25 kb unrearranged, deleted and inverted alleles. The same FAM probe (green) and reverse primer (gray) were used for all amplicons. Brown region indicates the 25 kb deletion or inversion.

**Fig. S2.**
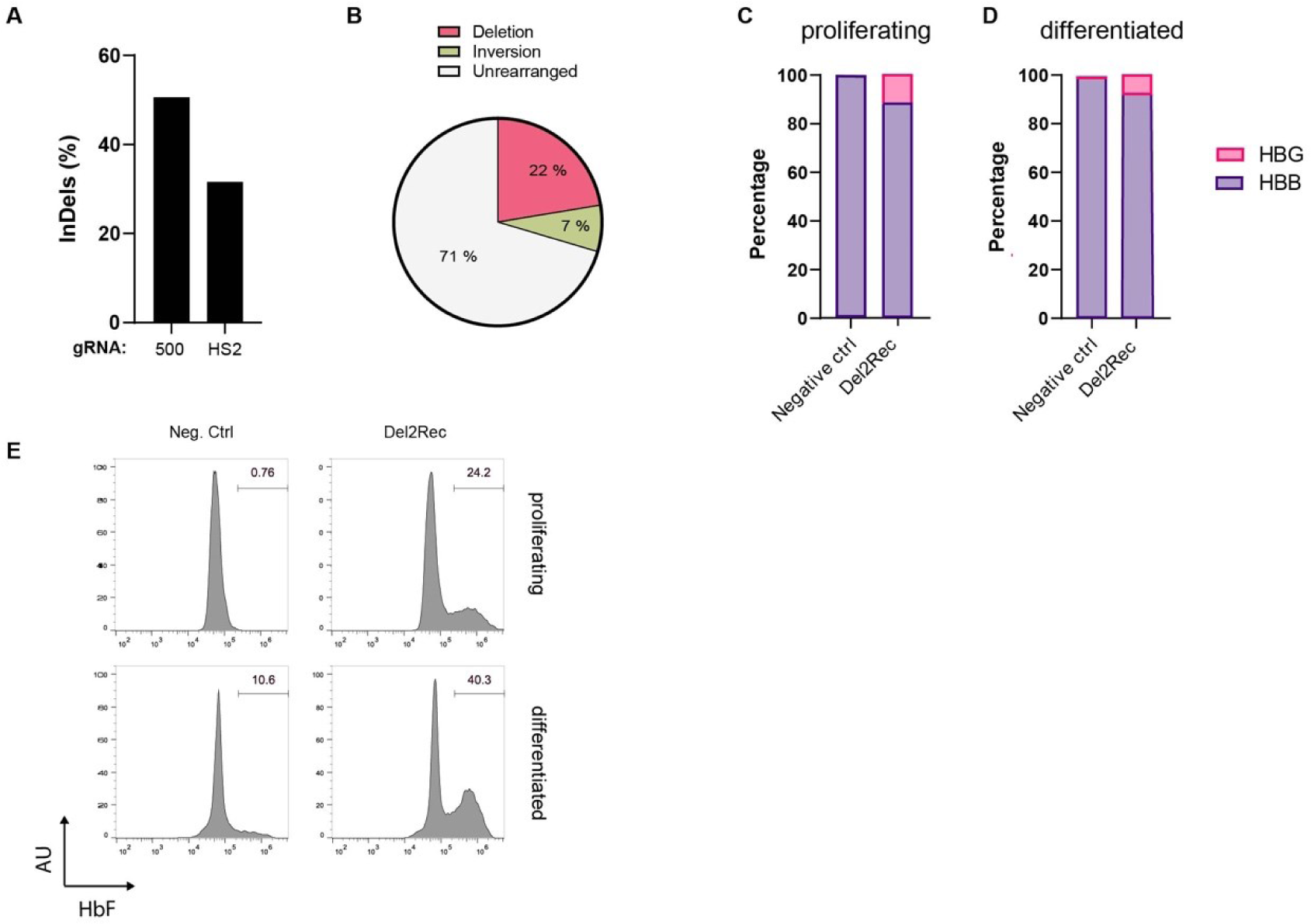
Linear recruitment of the LCR reactivates fetal globin in HUDEP-2 cells replicate experiment. a) Frequency of InDels, measured by TIDE analysis for 500 and HS2 gRNAs in single-gRNA transfected samples. b) Frequency of the 25-kb deletion (raspberry), inversion (olive) and unmodified (light gray), measured by ddPCR on Del2Rec transfected cells. c) RT-qPCR analysis of HBB and HBG mRNA levels in proliferating cells. HBB and HBG mRNA expression was normalized to actin mRNA. Displayed as a percentage of total HBB+HBG mRNA. d) Same as c) but on 10 days differentiated cells. e) Flow cytometry plots showing the percentage of HbF-positive cells in proliferating cells (top row) and differentiating cells at day 10 (bottom row).

**Fig. S3.**
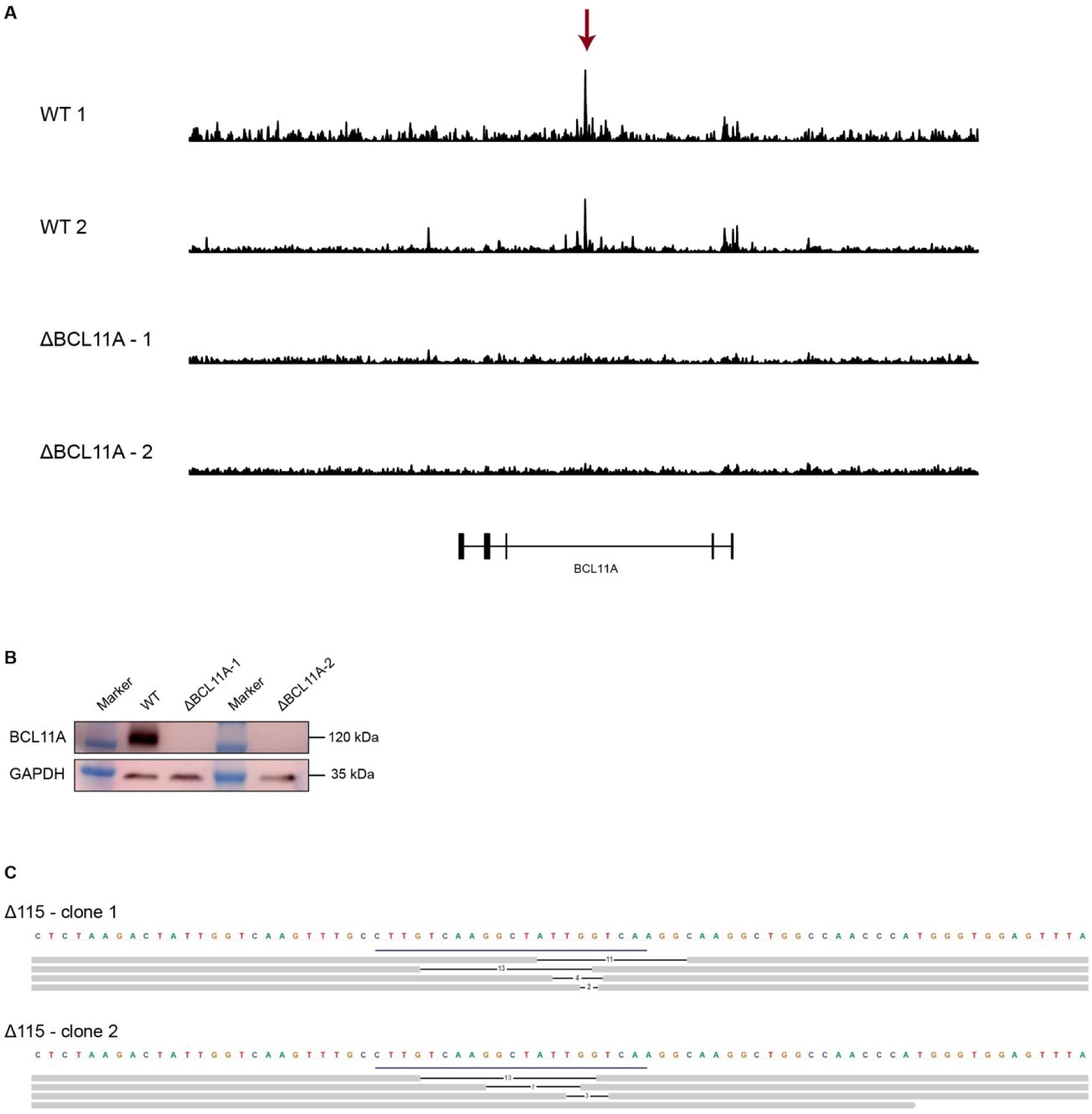
Analysis of Δ*BCL11A* and Δ115 HUDEP-2 clones. a) ATAC tracks of the *BCL11A* gene of WT and two **Δ***BCL11A* clones. Red arrow marks enhancer that is target by gRNA1617. Signal is scaled to genome wide TSS signal. b) Western Blot showing a WT and two Δ*BCL11A* proliferating clones, analyzed for BCL11A protein with GAPDH as loading control. c) Representative reads of ATAC-seq data of Δ115 clones showing the genotype of the two *HBG* promoters of two alleles. Clone 1 has 4 different deletions at the 115 site. Clone 2 has 3 different deletions and one WT 115 site. ATAC reads were aligned to *HBG1/2* promoters, reads spanning the 115 gRNA cute site were analyzed to determine genotypes.

**Fig. S4.**
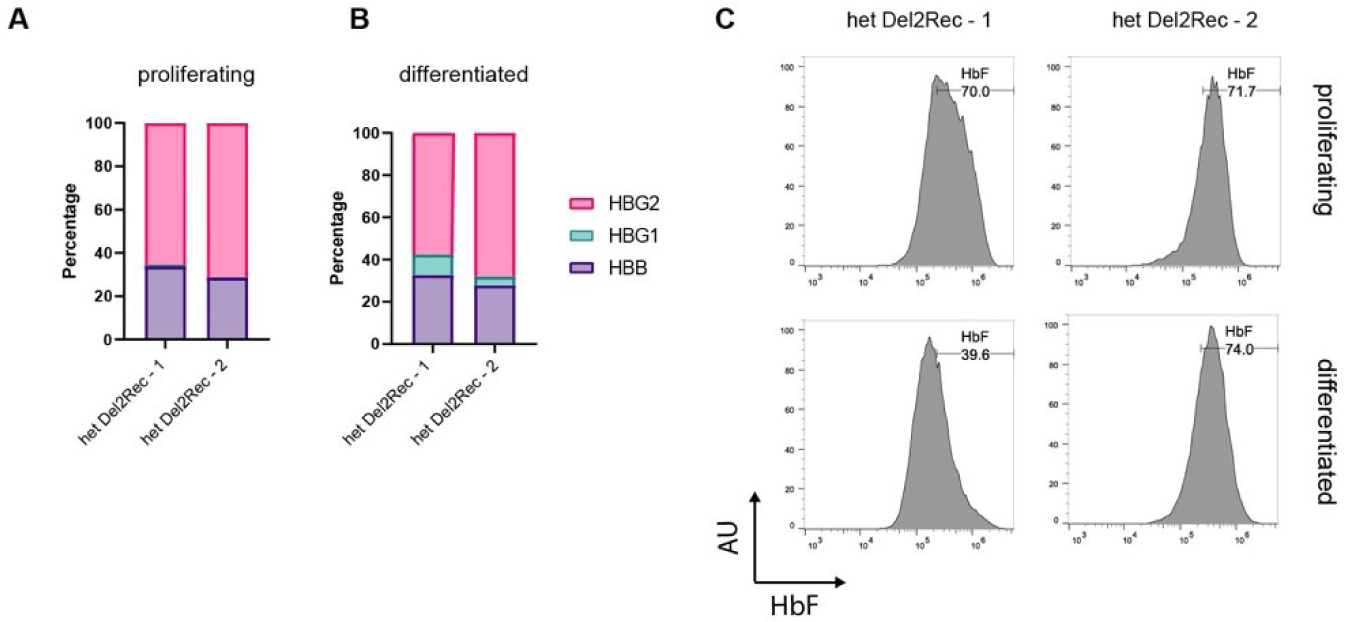
Heterozygous Del2Rec HUDEP-2 clones show high *HBG* reactivation. a) RT-qPCR analysis of HBB and HBG mRNA levels in proliferating clonal cell lines. HBB and HBG mRNA expression was normalized to actin mRNA. Displayed as a percentage of HBB+HBG1+HBG2 mRNA. b) Same as in a) but for cells differentiated for 10 days. c) Representative flow cytometry plots showing the percentage of HbF-positive cells in proliferating clonal cell lines (top row) and after differentiation (day 10, bottom row).

**Fig. S5.**
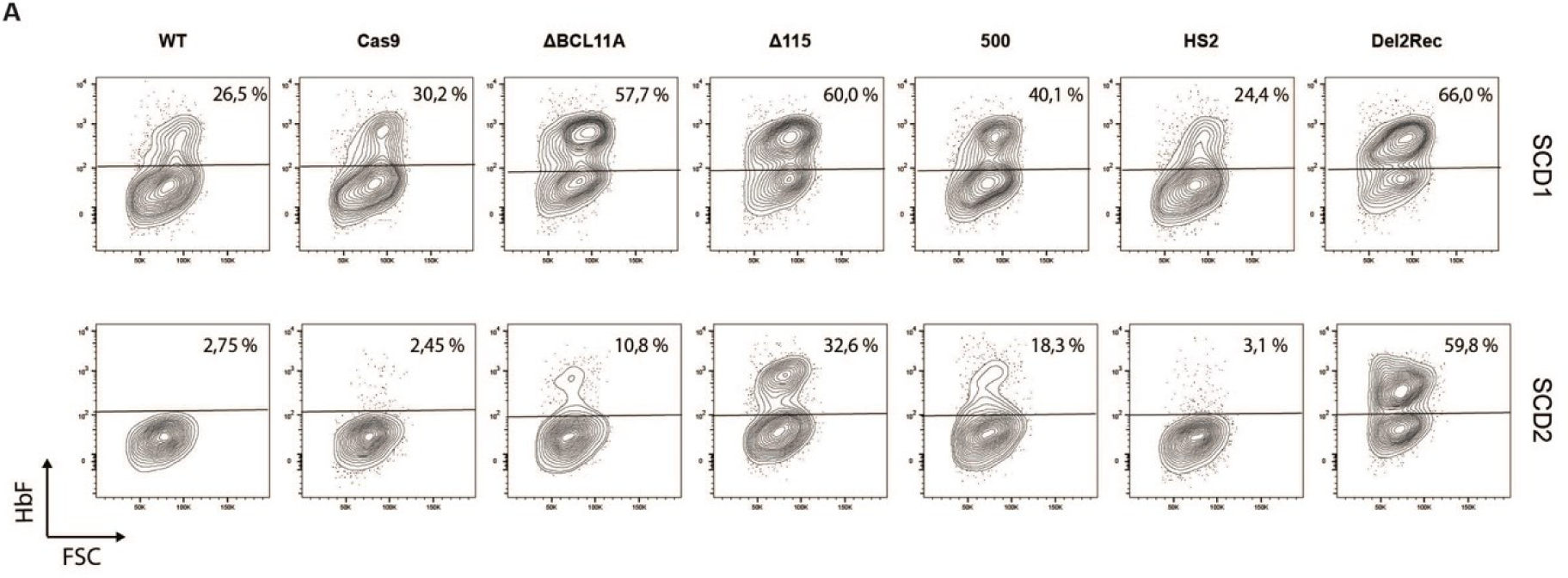
Linear recruitment of the LCR reactivates fetal globin in immortalized, patient-derived sickle cell disease cell lines. a) Flow cytometry plots showing the percentage of HbF positive cells in two REB-SCD cell lines. Note that SCD1 shows relatively high baseline HbF levels.

**Fig. S6.**
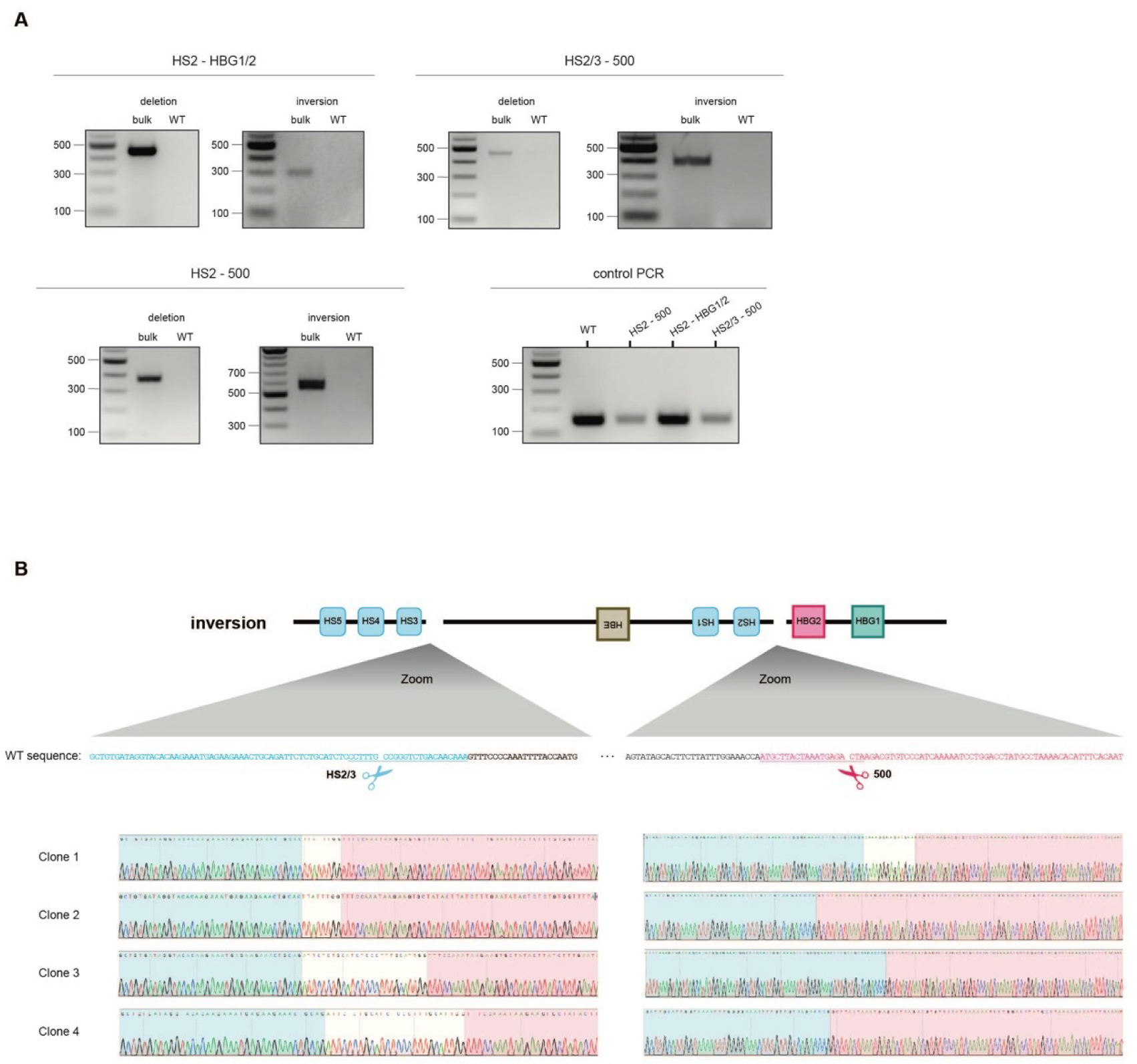
Bulk and clonal genotypes of cells treated with gRNAs *HBG1/2*-HS2 and 500-HS2/3. a) Genotyping PCR for deletion and inversion of bulk populations of conditions Cas9 only, gRNAs 500-HS2, gRNAs 500-HS2/3 and gRNAs *HBG1/2* – HS2 b) Sanger sequencing tracks of four HUDEP-2 inversion clones

**Fig. S7.**
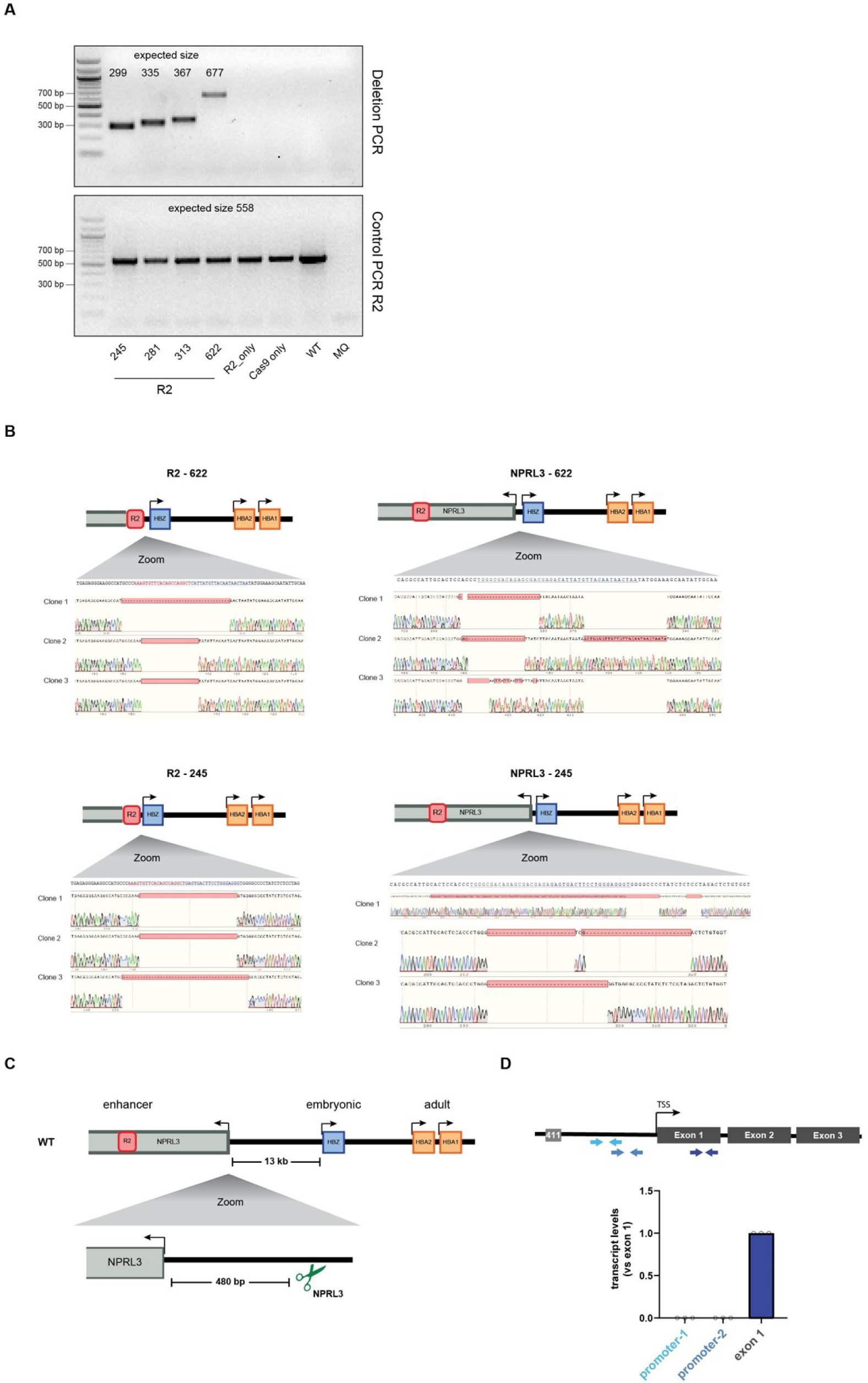
Bulk and clonal genotypes of cells treated with gRNAs *HBG1/2*-HS2 and 500-HS2/3. a) Genotyping PCR for deletions of bulk transfected HUDEP-2 cells, recruiting R2 to different distances from the *HBZ* TSS. b) Sanger sequencing tracks of HUDEP-2 clones containing a deletion from R2 (left) or NPRL3 (right) to the *HBZ* promoter, using a gRNA at 622 (top) or 245 (bottom) bp from the *HBZ* TSS. c) Schematic overview of clonal cell lines in which the *HBZ* promoter was recruited to the *NPRL3* promoter using a gRNA 490 bp to the right of the *NPRL3* TSS. d) Schematic representation of primers used in RT-qPCR analysis of HBZ transcripts (top). Fold-change normalized for primer efficiency based on DNA amplification with *HBZ* exon 1 set to 1, shown in arbitrary units (bottom). Each dot represents a clone, bar plot represent average.

**Fig. S8.**
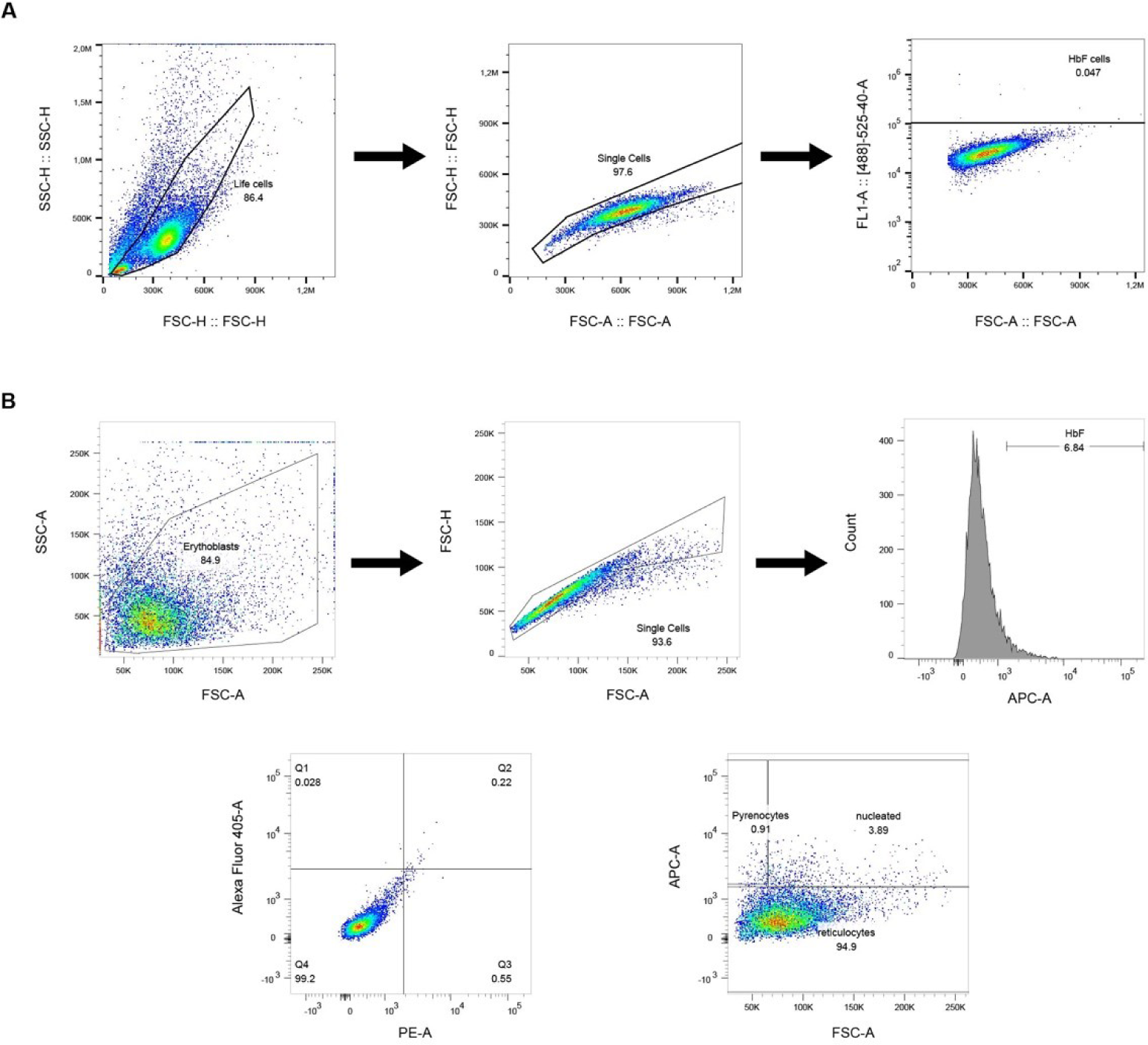
Representative flow cytometry plots from HUDEP-2 cell lines and HSPCs illustrating the gating strategy. a) HUDEP-2 were first gated on a forward scatter (FSC-H) versus side scatter (SSC-H) plot. Doublet exclusion was performed by selecting single cells based on FSC-H plotted versus FSC-A (FSC-area). HbF-positive gate was set by gating out the unstained sample. b) HSPCs were first gated on a forward scatter versus side scatter plot. Doublet exclusion was performed by selecting single cells based on FSC-H plotted versus FSC-A. HbF-positive gate was set by gating out the isotype stained sample.

## Materials and Methods

### HUDEP-2 cell line

HUDEP-2 cells were provided under an MTA by the RIKEN BRC through the National BioResource Project of the MEXT / AMED, Japan. Cells were cultured in Cellquin proliferation medium, which consists of fully defined serum free Iscove’s Modified Dulbecco’s Medium (IMDM), supplemented with human holotransferrin (300 μg/ml), Insulin (10 μg/ml), Na-pyruvate (100 □M), Penicillin/Streptomycin (P/S, 100 U/ml), L-glutamine (2 mM), lipid mix, Poly(vinyl alcohol) (PVA) (1 g/L), EPO (2 U/μL), dexamethasone (1 μM), doxycycline (3-6 μg/mL), and human recombinant stem cell factor (hSCF, 100ng/ml) (see Table S2) (20). To differentiate HUDEP-2 cells medium was switched to differentiation medium (see Table S2).

After 7 days, doxycycline was removed from the media and the cells were cultured for an additional 3 days.

### SCD cell lines

The Rotterdam erythroblast (REB-SCD) cell lines (REB-SCD46 (SCD1) and REB-SCD34 (SCD2)) have been generated from PBMCs obtained from patients with SCD (Majied and Philipsen, manuscript in preparation) using doxycycline inducible expression of the HPV-E6/E7 oncogenes as described previously to generate the HUDEP-2 cell line(21). In the proliferation medium 2 mg/L doxycycline was added to the medium to induce HPV-E6/E7 expression to immortalize the cell lines. The cells were cultured as described(20), in a fully defined serum free Iscove’s Modified Dulbecco’s Medium (IMDM) supplemented with cytokines, human holotransferrin (300 ug/ml), Insulin (10 ug/ml), Na-pyruvate (100 uM), Penicillin/Streptomycin (50 mg/L), L-glutamine (2 mM), Cholesterol (2.5 mg/L), L-a-phosphatydilcholine (2.5 mg/L), Oleic Acid (1.5 mg/L), Poly(vinyl alcohol) (PVA) (1 g/L), 2 U/μL EPO, 1 μM dexamethasone, 3-6 μg/mL doxycycline, and human recombinant stem cell factor (100ng/ml) (see Table S3). The cultures were maintained at 37 °C with 5% CO2 at a cell density of 0.2-1.0×10^6^ cells/ml. The medium was refreshed every second day. Cell lines were differentiated for 4 days in IMDM medium supplemented with 3% Omniplasma, EPO (10 U/mL), transferrin (330 μg/ml) and heparin (3 U/mL). At day two of differentiation the medium was refreshed.

### CD34+ HSPC isolation and Primary cell cultures

Human material was obtained after informed consent, mobilized peripheral blood (MPB) was obtained from leukapheresis material (Sanquin, The Netherlands). CD34+ cells were isolated from fresh MPBs using a Myltenyi CD34 kit and cryopreserved using IMDM + 20% FCS and 5% final DMSO concentration. Purity was determined using flowcytometry using the antibodies, listen in Table S4. All donors used had over 95% of CD34+ HSPCs.

Cryopreserved CD34^+^ HSPCs were thawed and cultured in IMDM media (PAN biotech, P04-20251K), supplemented with a cytokine maintenance cocktail containing stem cell factor (SCF) (100 ng/mL), Fms Related Receptor Tyrosine Kinase 3 ligand (FLT3LG) (100 ng/mL), and thrombopoietin (TPO) (10 ng/mL). All cells were precultured for 16–30 hours before nucleofection with Cas9 RNPs.

Directly after nucleofection, room temperature culture medium was added and cells were spun to remove residual nucleofection buffer. Cells were cultured with IMDM media including interleukin (IL) 3 (100 ng/mL), IL6 (100 ng/mL), TPO (10 ng/mL), and SCF (100 ng/mL) at 37°C, 5% CO_2_ for 4-5 consecutive days. The cells were further cultured to pure pro-erythroblasts for 4-10 days in IMDM (PAN biotech) supplemented with (1U/ml Epo), 100ng/ml SCF, 1µM dexamethasone (a glucocorticoid agonist) and 333 µg/ml holotransferrin(22). Differentiation of established pro-erythroblast cultures was initiated by culture in IMDM media (PAN biotech) supplemented with 5% human plasma, 10U/ml Epo and 1mg/ml holotransferrin(22). Cell counts were measured in duplicate on an Innovatis Casy TTC Cell Counter & Analyser System (Roche, 62687).

### RNP incubation and nucleofection in primary cells

For gRNA screening the ALTR crRNA system was used (IDT). Equal amounts of crRNA and Tracer RNA were incubated for 5 minutes at 95°C, followed by benchtop cooling. For other experiments gRNAs were used from IDT. RNPs were formed by incubating 9 µg Cas9-3xNLS, purified as described before(23), with 200 pmol of gRNA or gRNA for 10 minutes at room temperature. CD34^+^ HSPCs were collected and resuspended in 20 µl nucleofection buffer from Amaxa (P3), mixed with RNPs and nucleofected using program EO100 as previously described(23) .

### CRISPR editing in HUDEP-2 and REB-SCD cell lines

gRNAs are listed in Table S1. Nucleofections were carried out with in-house purified 3xNLS-Cas9(24), using the AmaxaTM P3 Primary Cell 4D-NucleofectorTM X Kit (EW-113). Single-cell clones were screened by PCR followed by Sanger sequencing confirmation. gRNA targeting AAVS1 was used as a negative control in bulk HUDEP-2 experiments.

### RT-qPCR

Cells were homogenized using TRIzol (Life Technologies). RNA was isolated either by phenol-chloroform extraction or using the Direct-tol^TM^ RNA MiniPrep Kit (Zymo, Cat No: R2052). cDNA was synthesized using M-MLV RT H (-) point enzyme (Promega). Gene expression was normalized to HBA or Actin B (ACTB). Primers for RT–qPCR are listed in Table S1.

### HbF staining and flow cytometry

Cells were fixed in 0.05% glutaraldehyde for 10 min at room temperature and then permeabilized with 0.1% Triton X-100 for 3 min. Cells were stained with HbF-FITC conjugated antibody (MHFH01, lot.2282927) for 15 min in the dark. Flow cytometry was performed on a Beckman Coulter Cytoflex S. Data were collected with CytExpert 2.3.1.22 and analyzed with FlowJo 10.8.0 software (Extended Data Figure 7).

### Lentivirus production and transduction

To generate inversions and *HBA* locus cell lines a founder cell line containing Cas9-Blast (addgene #52962) was used. Cells were maintained in medium containing 1 μg/ml blasticidine (InvivoGen, ant-bl-05). sgRNA sequences (Table S1) were cloned into a modified version of pU6-gRNA EF1 Alpha-puro-T2A-GFP or BFP lentiviral targeting plasmid (addgene #60955). The medium was refreshed 24 h after transfection. Medium containing the virus particles was harvested 48h after transfection by passing through a 0.45 m filter and concentrating it 8x using Amicon Ultra centrifugal filters, 50kDa MWCO (UFC905008). This was added to the HUDEP-2 cells in Cellquin medium supplemented with 6 g/mL polybrene (Merck) and 1% HEPES. Cells were centrifuged for 90 min at 800 g at RT. The cells were refreshed 24h after transduction.

### Western blots

Antibodies are listed in Table S4.

Whole cell lysates were run on polyacrylamide gel and immunoblotted using standard procedures.

### HPLC

1×10^7^ cells were collected and analyzed for hemoglobin expression by high-performance cation-exchange liquid chromatography (HPLC) on Waters Alliance 2690 equipment as described previously(25).

### ddPCR

Primers and probes are listed in Table S1.

Reaction mixtures of 20 mL volume were prepared containing 1xddPCR Master Mix (Bio-Rad, Hercules, CA, cat#12005909), relevant primers and probe (400 nM primers and 100 nM probe), TERT reference (dHsaCP2500351), and 3 µL of MseI digested gDNA (20 ng for HUDEP-2, REB-SCD and for primary cells). Droplet generation was performed as described(26). The droplet emulsion was transferred to a 96-well propylene plate (Eppendorf, Hamburg, Germany) and amplified in a conventional thermal cycler (T100 thermal cycler, Bio-Rad, Hercules, CA). Thermal cycling conditions consisted of 95C for 10 min; 94C for 30 s, 60C for 1 min, and 72C for 2 min (40 cycles); 98C for 10 min (1 cycle); and 12C hold. After PCR, the 96-well plate was transferred to a droplet reader (Bio-Rad, Hercules, CA). Acquisition and analysis of the ddPCR data were performed with the QuantaSoft. Relative frequencies of each rearrangement (deletion, inversion or unrearranged) was calculated as a fraction of the total (deletion + inversion + unrearranged).

### ATAC-seq

ATAC-seq was performed using the Omni-ATAC protocol. Briefly, 200,000 cells were lysed with 0.1% NP-40, 0.1% Tween-20 and 0.01% digitonin, and incubated with in-house produced Tagment DNA Enzyme for 30 min at 37 °C. DNA was purified with QIAGEN MinElute Reaction Cleanup Kit. Library fragments were amplified using Phusion High-Fidelity PCR Master Mix with HF Buffer (Thermo Fisher Scientific, catalog no. F531S) and customized primers with unique single or dual indexes. Libraries were purified using AMPure XP beads (Beckman Coulter, catalog no. A63881) according to the manufacturer’s instructions. Libraries were evaluated using the Agilent Bioanalyzer 2100 with the DNA 7500 kit (catalog no. 5067-1504). Libraries were then pooled and sequenced in single (SE) or paired-end (PE) mode using a P2 flow cell on the NextSeq 2000 using Illumina-supplied kits as appropriate. Reads with trimmed with cutadapt 4.4 with --minimum-length 10 and removal of adapters, aligned with bowtie2 with parameters --local --very-sensitive-local --no-unal --no-mixed --no-discordant -- dovetail -X 1000. Non-primary alignments, not-properly paired reads (in case of PE reads) and alignments with mapping quality lower than 15 were filtered out with samtools 1.17 (view - F 260) and blacklisted against ENCODE Unified GRCh38 Blacklist ENCFF356LFX (https://www.encodeproject.org/files/ENCFF356LFX/) with bedtools 2.31.0. Duplications were retained because the single-end 51 bp reads have a higher likelihood of being duplicated while originating from separate fragments.

For visualization, bigwigs were created according to the ENCODE atac-seq-pipeline (https://github.com/ENCODE-DCC/atac-seq-pipeline). First we converted the BAM files to tagAlign BED files with bamtobed (bedtools 2.31.1) and created treatment bedgraphs during macs2 (2.2.9.1) peak calling, with artificial reads centralized 75bp up- and downstream of Tn5 transposase cutting sites, with parameters -f BED -g hs -p 0.01 --shift -75 --extsize 150 -- nomodel --bdg --SPMR --keep-dup all. Finally treatment bedgraphs were converted to bigwigs with bedGraphToBigWig (ucsc tools 3.3.5).

Bigwigs were displayed using rtracklayer 1.62.2 and GenomicRanges 1.54.1 packages. Bigwigs for clones being biological replicates were averaged with bigwigAverage (deeptools=3.5.2) and the final signal track was normalized against the average signal around transcription start sites (500bp up- and downstream).

### CUT&RUN

CUT&RUN was performed using the CUT&RUN Kit from Cell Signaling (catalog no. 86652S) according to the manufacturer’s instructions. Briefly, 0.5 million cells were harvested and washed twice in 1× wash buffer at room temperature and bound to 15 μl of Concanavalin A beads. Primary antibody incubation was performed in PCR tubes for 2 h at 4 °C with rotation.

H3K4me3 antibody supplied by the kit was used (1:50). After two washes, pAG-MNase was added and incubated for 1 h at 4 °C with rotation. After three washes, cells were resuspended in 1× wash buffer and pAG-MNase digestion was activated by adding 2 mM CaCl2. Samples were incubated on ice for 45 min, then 2× stop buffer was added to end the reaction. The samples were incubated at 37 °C for 10 min to release the captured chromatin fragments. The sequencing libraries were prepared using the NEBNext Ultra II DNA Library (New England Biolabs, catalog no. E7645) with 15–30 ng of input DNA. Sequencing libraries were pooled and paired-end (2 x 50 bp) sequenced on an Illumina Nextseq 2000 platform.

## Acknowledgements

We thank the members of our labs for discussions and feedback. We thank M. Bauer for help with setting-up ATAC-seq, C.Valdes-Quezada for input on the ddPCR and A. Didriksen for providing Cas9 protein. We thank the protein facility of the Netherlands Cancer Institute for providing the Tn5 protein. We acknowledge the Utrecht Sequencing Facility (USEQ) for providing sequencing service and data. USEQ is subsidized by the University Medical Center Utrecht and The Netherlands X-omics Initiative (NWO project 184.034.019). Research in the laboratory of WdL was financially supported by the EU Horizon 2020-funded Innovative Training Network ‘Molecular Basis of Human Enhanceropathies’ (Enhpathy,www.enhpathy.eu), under Marie Sklodowska-Curie grant agreement no. 860002, a NWO Groot grant (2019.012) from the Netherlands Organisation for Scientific Research (NWO) and Oncode Institute Base Funding. Work in the laboratory of SP was supported by TKI Health Holland (EMCLSH20006 and EMCLSH20025), ZonMw PSIDER consortium TRACER (10250022110001), EU Horizon Europe Pathfinder EdiGenT (101070903) and NWO Applied and Engineering Sciences Open Technology Programme (18947). Worked by the laboratory of EvdA was supported by Sanquin research fund l2842 and Sanquin Blood Supply grant PPODR21-07.

## Author contributions

A-KF, PHLK and WdL conceived and supervised the study. R. Majied, TV and SP designed and executed the experiments in SCD cell lines, HJMPV and EvdA designed and executed the experiments in CD34+ HSPCs and performed HPLC analysis, SJDT designed and performed the HBA experiments, with help from A-KF and MJAMV. A-KF performed experiments in HUDEP-2 cells, with help from MJAMV and R. Mohnani. A-KF performed the TIDE and ddPCR analyses, RG analyzed Cut&Run and ATAC-seq data. A-KF and WdL wrote the manuscript, with help from SJDT, R. Majied, SP, HJMPV and EvdA.

## Data availability

H3K4me3 CUT&RUN and ATAC-seq datasets are available at the Gene Expression Omnibus database under accession code GSE274030 and GSE274029, respectively.

